# Disruptions in cell fate decisions and transformed enteroendocrine cells drive intestinal tumorigenesis in Drosophila

**DOI:** 10.1101/2022.09.25.509404

**Authors:** Maria Quintero, Erdem Bangi

## Abstract

Most epithelial tissues are maintained by stem cells that produce the different cell lineages required for proper tissue function. Constant communication between different cell types that make up a tissue is essential to ensure that all cell lineages are produced at appropriate numbers and to mount regenerative responses to injury, infection, and other environmental stresses. Cancer-driving alterations change the intrinsic properties of transformed cells and disrupt stem cell regulation, cell fate decisions, and cell-cell communication within transformed tissue. However, mechanisms by which these processes are disrupted and co-opted to support tumorigenesis are not well understood. Here, we report a novel genetic platform, PromoterSwitch, that allows targeting of genetic manipulations to a small subset of cells of any tissue or cell type of interest and all their subsequent progeny. We use this technology to generate large, transformed clones derived from individual stem/progenitor cells in the adult Drosophila intestine. We show that cancer-driving genetic alterations representing common colon tumor genome landscapes drive disruptions in cell fate decisions within transformed clones and changes in the relative abundance of different intestinal cell lineages. We also uncover a critical, context-dependent role for the differentiated, hormone-producing enteroendocrine (EE) cells in the growth and maintenance of transformed clones. Our analysis in different genetic contexts provides insights into how the intrinsic properties of transformed cells —dictated by the genetic alterations they carry— determine their response to their environment and dependence on niche signals. A better mechanistic understanding of disruptions of cell-cell communication, stem cell regulation, and cell fate decisions within tumors could reveal novel vulnerabilities and druggable regulatory nodes that can be exploited for therapy. Understanding how tissues respond to the emergence of cells with cancer-driving genetic alterations also provides insights into stem cell biology and epithelial homeostasis.

## INTRODUCTION

Precise regulation of stem cell behavior and cell fate decisions is critical to maintaining tissue integrity and homeostasis. Stem cell proliferation, self-renewal, and differentiation are coordinated by a host of intrinsic, lineage-specific factors and extrinsic signals that mediate communication among different cells that make up a tissue and their microenvironment^1^. These interactions ensure that all cell lineages essential for proper tissue function are produced and maintained at appropriate numbers. Disruption of these cell-cell communication mechanisms can alter the relative abundance of different cell lineages within a tissue, resulting in tissue dysfunction, overproliferation, and cancer. It can also interfere with the ability of a tissue to appropriately respond to injury, infection, environmental stresses, and the emergence of cells with cancer-driving genomic alterations.

Most solid tumors, including intestinal tumors, are composed of cell types typically found in their tissue of origin^2^. Cancer-driving genetic alterations that accumulate in tumors cooperate to disrupt normal stem cell proliferation and differentiation mechanisms to establish tumors composed of cells with different intrinsic properties than their wild-type counterparts^3^. In addition, homeostatic cell-cell communication mechanisms that ensure tissue integrity and function are disrupted and co-opted to support the growth and maintenance of transformed tissue^4,5^. Molecular mechanisms underlying interactions of different transformed cell lineages, their wildtype neighbors, and their contributions to tumorigenesis are not well understood.

Most tumors carry concurrent alterations in multiple genes, and there is extensive genetic heterogeneity among different tumors of the same type^6–9^. As a result, the specific genomic landscape of a tumor cell will determine not only its intrinsic properties and behavior but also the extracellular signals it produces to communicate with its neighbors and how it responds to its environment. Therefore, a comprehensive understanding of how homeostatic mechanisms are altered to support tumor development requires studying cell fate decisions and cell-cell interactions in experimental systems that reflect the genetic complexity and diversity of human tumor genome landscapes.

The intestinal epithelium is one of the most rapidly renewing tissues in the body and serves as an attractive and relevant paradigm for studying stem cell regulation and epithelial homeostasis^10,11^. Both the fly and mammalian intestine are highly regenerative organs with stem and progenitor cells that produce two broad categories of differentiated cell types: absorptive enterocytes (ECs) and hormone-producing enteroendocrine (EE) cells^12–16^. Single-cell RNA sequencing analyses demonstrate that cell type-specific transcriptional signatures between Drosophila and mammalian intestinal cell types are evolutionarily conserved^17,18^. Signaling pathways governing intestinal homeostasis, stem cell renewal, differentiation, and interactions between the intestinal epithelium and its microenvironment are also highly conserved in Drosophila^19–26^. These similarities provide confidence that the behavior of the Drosophila intestinal cell types reflects those of the mammalian intestine.

Intestinal cells use several highly conserved signaling pathways to communicate with each other to ensure tissue integrity and proper tissue function^27,28^. The best characterized cell-cell interactions include signals sent from differentiated cells to intestinal stem cells (ISCs) to regulate stem cell proliferation and differentiation. For instance, JAK/STAT pathway ligands (upd1-3) are upregulated in ECs upon injury and signal ISCs to promote their proliferation^29^. Similarly, neighboring cells express and secrete multiple EGFR ligands to stimulate ISC proliferation in response to infection and injury^30^. There is evidence for both mechanisms playing a role under normal homeostatic conditions as well^30,31^. Enteroendocrine cells also send signals to other cell types, including ISCs, directly and indirectly^32–34^. For instance, the peptide hormone Tachykinin (Tk) signals the overlying muscle to produce and secrete DILP3, which in turn stimulates ISC proliferation through the Insulin Receptor (InR)^32^. Intestinal stem cell tumors in Drosophila depend on secreted signals from wildtype ECs and non-autonomous JNK and Hippo signaling^19,35–38^, but a role for the EE lineage in this context has not been established. In addition, human tumors typically have high numbers of transformed differentiated cells, which may also secrete these niche signals, reducing tumors’ dependence on non-autonomous signals from the surrounding epithelium. Relative contributions of different transformed versus wild-type cell lineages during tumorigenesis remain largely unknown.

We previously leveraged fundamental similarities between the mammalian and Drosophila intestine^28,39,40^, and the Gal4/UAS/Gal80(ts) system, a repressible targeted expression system for tissue-specific genetic manipulations in Drosophila^41,42^, to study intestinal transformation by genetically manipulating Drosophila orthologs of genes recurrently mutated in colon tumors^43–47^. These models focused on five genes, APC, KRAS, TP53, SMAD4, and PTEN, which lead to deregulation of Wnt, RAS/MAPK, DNA damage response/apoptosis, TGF-β, and PI3K pathways, respectively^48,49^. They captured critical aspects of tumorigenesis and showed that many tumor phenotypes are emergent features of interactions between multiple transgenes^50^,^51^. We also established and drug-screened more personalized multigenic models based on patients’ specific tumor genome profiles and demonstrated that Drosophila models could identify drug combinations effective in cancer patients^52,53^.

Exploring cell-cell interactions and fate decisions during intestinal transformation requires studying discrete clones originating from individual cells of defined lineages and capturing the cell lineage heterogeneity typically found in colon tumors. Targeting cancer-driving multigenic combinations to different cell lineages using cell-type specific Gal4 lines results in the transformation of many cells in the intestine, rapidly compromising tissue integrity. Furthermore, too many clones in close proximity make it challenging to identify discrete clones and study cell-cell interactions at the clone boundary. Lastly, cell-type specific promoters mediating transgene expression can become silenced as transformed cells attempt to differentiate.

To overcome these challenges, we established a new genetic platform, PromotorSwitch (PS), inspired by the CoinFLP method^54^. Using this design, we could generate and study a small number of clones derived from individual cells of defined cell lineages. Targeting our multigenic models to a small number of individual stem/progenitor cells in the adult intestine results in large multicellular clones that show multiple key indicators of oncogenic transformation. They contain all cell lineages typically found in the normal intestine, albeit at higher numbers, and cells that co-express multiple cell lineage markers, reflecting the cell lineage heterogeneity observed in epithelial tumors. We also find genotype-dependent differences in the relative abundance of different intestinal cell lineages. Finally, we identify a tumor-promoting role for the transformed EE cells during intestinal tumorigenesis and describe an underlying molecular mechanism. These findings provide insights into cell fate decisions and cell-cell communication during tumor progression in a whole-tissue context and offer a possible mechanism for a clinical observation that colon tumors with high numbers of EEs appear to be more aggressive with a worse prognosis^55,56^.

## RESULTS

### Generating Transformed Clones Using the PromoterSwitch Design

Here, we introduce PromoterSwitch, an intersectional system that combines UAS/Gal4/Gal80(ts) and the site-directed FLP/FRT recombination methods (Figure 1a,b). In this design, the expression of FLP recombinase, which can excise any DNA sequence flanked by its FLP Recognition Target (FRT) sites^57^, is first initiated using a cell-type specific Gal4. Then, most initially targeted cells undergo a FLP recombinase-mediated switch that permanently shuts down transgene expression, while a small subset and their subsequent progeny permanently lock in transgene expression by switching to a ubiquitous promoter to regulate Gal4 expression. As this “promoter switch” only occurs in initially targeted cells, cell specificity of genetic manipulations is preserved. To simplify the genetic background of our experimental animals and allow the introduction of additional transgenes for mechanistic studies, we consolidated the three transgenes required for the PS design into a single construct using our multigenic vector platform^52,53^ (Figure 1a).

**Figure 1.**
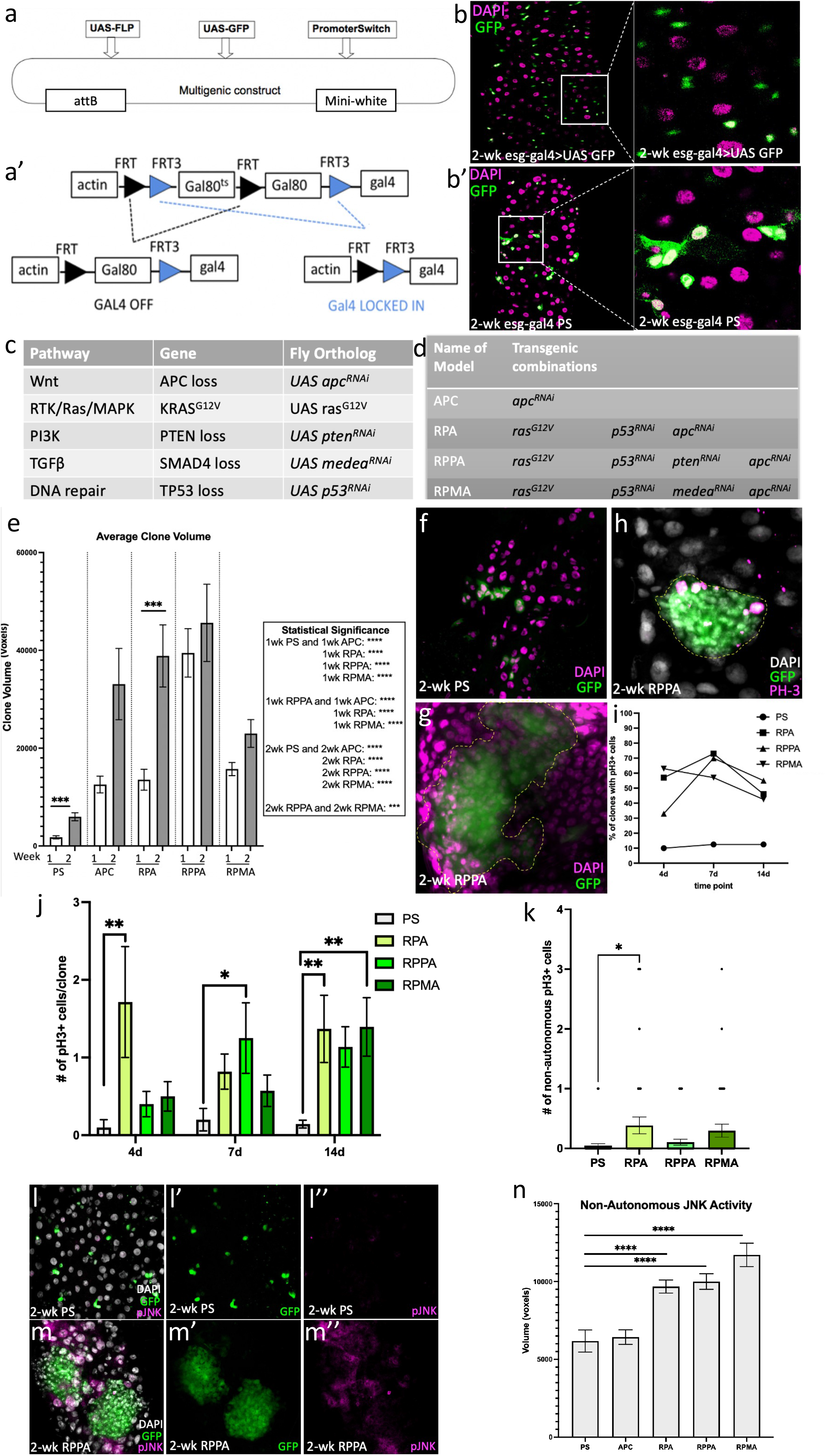
Generating transformed clones in the adult Drosophila intestine using the PromoterSwitch (PS) design. **(a)** PS multigenic construct contains Gal4-inducible *UAS-FLP* and *UAS-GFP* transgenes and the *PS* transgene **(a’**). PS construct is activated by crossing it to a cell-type specific Gal4 and inactivating Gal80^ts^ with a temperature shift. Once induced from the *UAS-FLP* transgene, FLP recombinase can either act on 1) the FRT pair (black triangles), excising the Gal80^ts^ coding sequence and resulting in ubiquitous Gal80 expression and inhibition of Gal4 activity, or 2) the FRT3 pair (blue triangles), excising both Gal80^ts^ and Gal80 coding sequences and resulting in ubiquitous expression of Gal4. **(b,b’)** Adult midguts two weeks after induction of GFP expression using *esg-gal4* only **(b)** and with PS **(b’)**. Panels on the right show magnified regions outlined by the white boxes in the left panels. PromoterSwitch allows targeting a small subset of *esg-gal4* expressing cells (green) and their subsequent progeny, resulting in small multicellular clones that include large differentiated cells. **(c)** Commonly deregulated pathways in human colon tumors, cancer driver genes selected to represent each pathway, and transgenes used to manipulate their Drosophila orthologs **(d)** Transgenic combinations used in this study. **(e)** Analysis of clone volume. Significant differences in clone volume among different genotypes are time points were observed (Multiple unpaired t-tests with Holm-Šídák correction) **(f,g).** Example clones used in volume analysis. Targeted cells in green and nuclei (DAPI) in magenta. **(h-k)** Proliferation analysis using the mitotic marker phospho-Histone 3. **(h)** Example clone used for quantification of proliferation. Clone in green, nuclei (DAPI) in gray, and pH3 in magenta. **(i)** Fraction of clones with pH3+ cells. **(j)** Number of pH3+ cells/clone, and **(k)** Number of pH3+ cells in close proximity to the clones. Significant differences in proliferation rates within and surrounding clones were observed (One-Way ANOVA). **(l,m)** An example two-week-old RPPA clone (green, **m)** showing both autonomous and non-autonomous JNK pathway activation (pJNK, in magenta). **(n)** Quantification of non-autonomous JNK activity surrounding clones (Multiple unpaired t-tests with Holm-Šídák correction). **(e, j, k,n)** Error bars represent standard error of the mean (SEM); *: P ≤0.05, **: P≤ 0.01, ***: P≤ 0.001, ****: P≤ 0.0001. Quantifications shown in **e** and **n** were performed using a custom pipeline in CellProfiler (see methods for details).

We then used the PS design and *esg-gal4*, expressed in intestinal stem and progenitor cells, to target our multigenic combinations to a small number of individual cells in the adult intestine (Figure 1c,d). Targeted cells overproliferated, resulting in large, multicellular clones compared to GFP-only control clones (Figure 1e-g). Consistent with these size differences, experimental clones had higher proliferation rates (Figure 1h-j). Of note, we also found higher levels of proliferation in wild-type cells close to experimental clones with specific genotypes (Figure 1k), suggesting a non-autonomous effect on proliferation. Genotype-dependent differences we observed in clone sizes, growth trajectories, and proliferation rates (figure 1e,i,j) highlight the importance of genetic context in determining the behavior of transformed cells.

Experimental clones with all genotypes also show activation of Src, JNK, MAPK, and AKT signaling pathways and induction of MMP1 expression —all critical indicators of oncogenic transformation (Figure 1l,m, Figure S1). We also observed non-autonomous activation of these markers in wild-type tissue surrounding transformed clones (Figure 1n, Figure S1), which is significant as these pathways have essential roles in intestinal homeostasis and tumorigenesis^10,19^. They are also critical regulators of cell competition and compensatory proliferation in different experimental contexts^58^. These models recapitulate various cancer hallmarks, such as increased proliferative signaling and activation of cancer-relevant pathways, providing an excellent model system to study cell fate decisions and the roles of different cell lineages in establishing and maintaining transformed clones.

### Cell lineage composition of transformed clones

Next, we analyzed the cell type composition of transformed clones using previously established markers for each lineage (Figure 2). At any given point, a typical control clone includes 1-2 ISCs, and may also include a small number of differentiated and progenitor cell lineages (0-2 each, Figure 2a,c,e,g,i,k,m,o). Transformed clones had all cell lineages typically found in the normal intestine (Figure 2a,d,e,h,i,l,m,p). Still, we noted differences in the relative abundance of some cell lineages across different genotypes and over time. The relative abundance of ISCs and enteroblasts (EBs), which are progenitors of absorptive enterocytes (ECs), in transformed clones were similar to controls (Figure 2a,e). However, RPA, RPPA, and RPMA clones all had a significantly higher relative abundance of EE cells (Figure 2i), and that of ECs was significantly lower in RPMA clones (Figure 2m). When we focused our analysis on large clones only, we found a significantly higher relative abundance of ISCs and EBs in large RPA and RPMA clones, respectively, indicating that these clones become less differentiated over time (Figure 2b,f). Consistent with this finding, smaller clones tended to have a higher fraction of differentiated cells than larger ones (Figure 2j,n). Of note, even in cases where the fraction of cells of a particular lineage is similar across genotypes (e.g., ISCs, Figure 2a), their number is much higher because of the larger overall size of transformed clones (Figure 1e).

**Figure 2:**
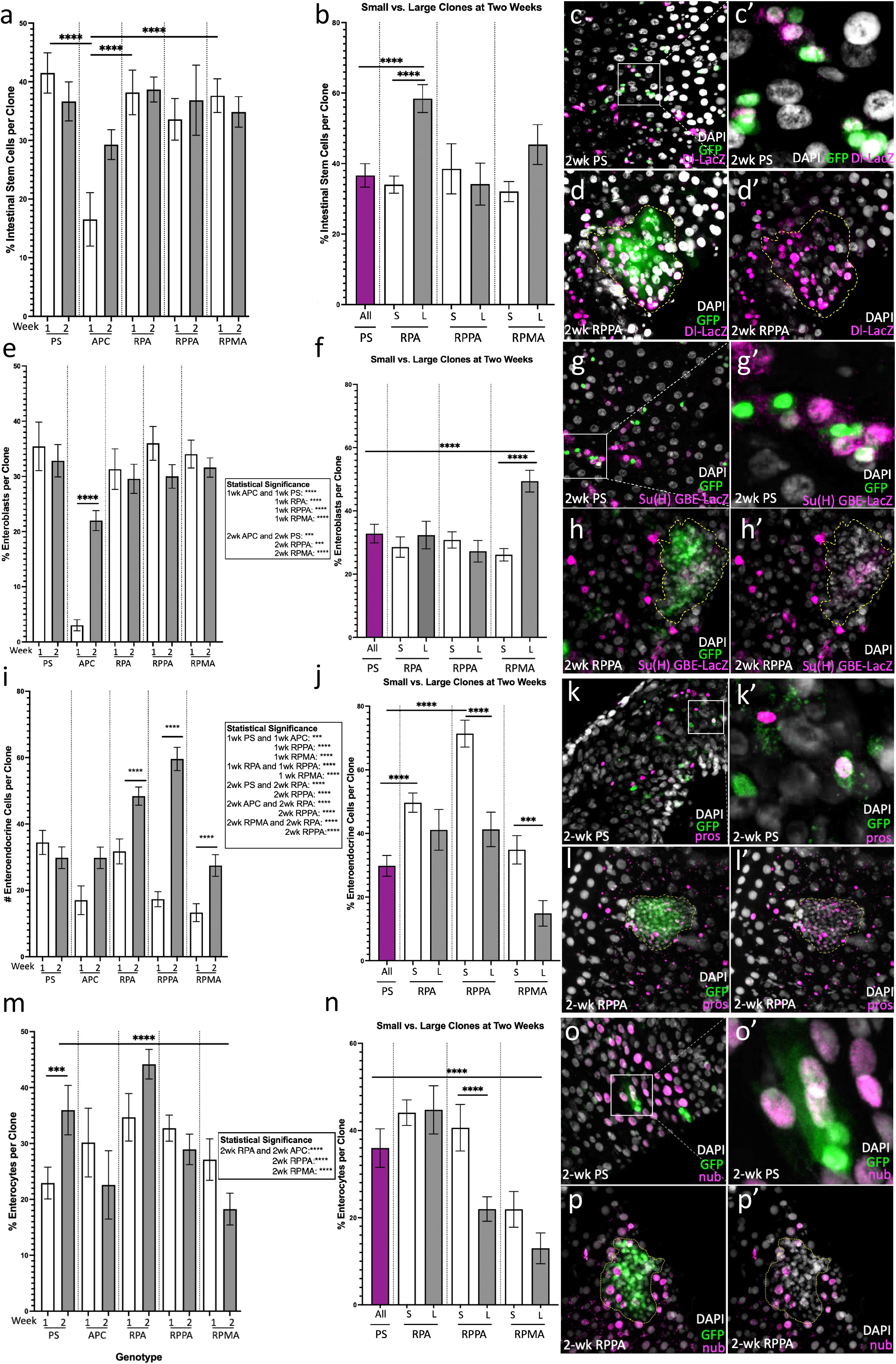
Analysis of cell type composition of transformed clones. **(a,e,i,m)** Fraction of cells positive for the ISC marker Dl-LacZ **(a)**, EB marker Su(H)-GBE-LacZ **(e)**, EE marker Prospero (pros, **i**), and EC marker Nubbin (nub, **m**) per clone at indicated time points and genotypes. **(b, f,j,n)** The fraction of cells positive for different lineage markers shown separately in two-week-old small (S) and large (L) experimental clones. PS alone produces GFP-only control clones that are all small. Example control **(c,g,k,o)** and experimental clones **(d,h,l,p)** used for cell lineage quantification. Clones are green, nuclei (DAPI) in gray, and cell lineage markers in magenta. **(c’,g’,k’,o’)** Magnified regions squared off in panels **(c,g,k,o)** focusing on GFP-only control clones. **(d’,h’,l’,p’)** Transformed clones shown in **(d,h,l,p)** with GFP removed and clones outlined in yellow dashed lines to allow better visualization of lineage markers. Quantifications were performed using a custom pipeline in CellProfiler (see methods for details). Error bars represent the standard error of the mean (SEM). *: P ≤0.05, **: P≤ 0.01, ***: P≤ 0.001, ****: P≤ 0.0001 (Multiple unpaired t-tests with Holm-Šídák correction, PRISM Software).

### Transformed clones contain cells that co-express multiple cell fate markers

Our analysis of cell type composition of transformed clones shows time, clone size, and genotype-dependent differences in the relative abundance of some cell lineages, indicating a disruption of stem cell regulation and cell fate decisions within transformed clones. To determine whether transformed clones also contain cells with abnormal cell fates, we conducted a colocalization analysis of cell lineage markers, focusing on combinations of differentiation and stem/progenitor markers. This analysis revealed a small number of cells that carry markers of multiple cell lineages in transformed clones. For instance, approximately 5% of large RPA clones contained cells positive for both Dl-LacZ and Pros, markers of ISC and EE cell fates, respectively, while no such cells were present in control clones (Figure 3a,b). We corroborated these findings using Tachykinin (Tk), a peptide hormone with an essential function in lipid metabolism and homeostasis secreted by a subset of EE cells^33^ (Figure 3c).

**Figure 3.**
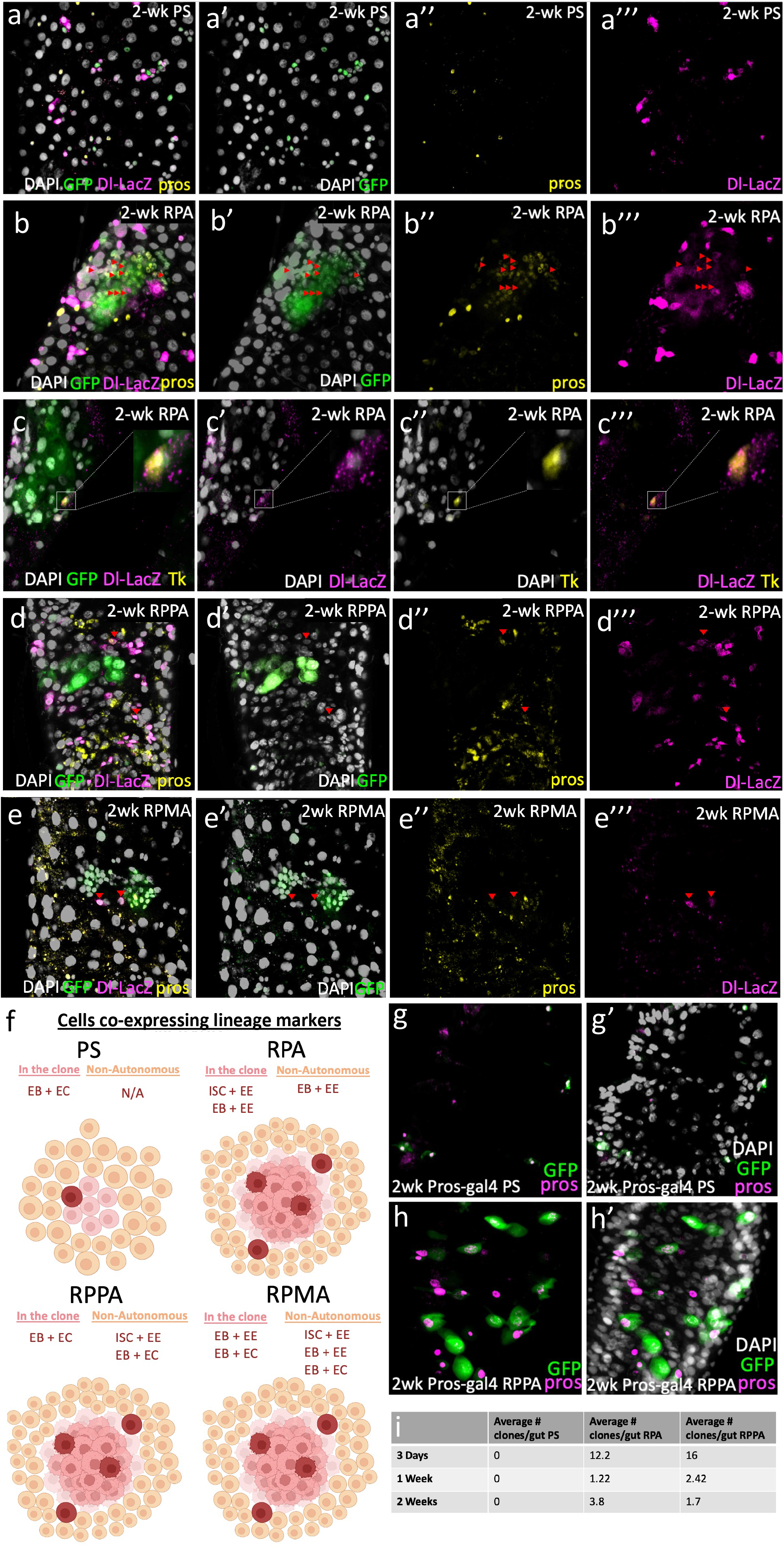
Transformed clones contain cells that co-express multiple cell fate markers. Intestines with example two-week-old GFP-only control **(a)**, RPA **(b,c)**, RPPA **(d),** and RPMA **(e)** clones immunostained for indicated markers. Clones are in green, nuclei (DAPI) in gray, Dl-LacZ in magenta, pros **(a,b,d,e),** and Tachykinin (Tk, **c**) in yellow. A cell co-expressing Dl-LacZ and Tk is outlined in a white rectangle and a magnified view shown in the small square to the right of each image in **(c)**. **(f)** A graphical representation of cells co-expressing multiple cell fate markers observed within and surrounding clones with different genotypes (made with BioRender). **(g,h)** Cells/clones (green) generated by targeting individual EE cells using *pros-gal4* and the PS construct. **(g)** GFP-only controls. **(h)** Small multicellular clones generated by RPPA. Nuclei (DAPI) are in gray, and pros in magenta. **(i)** The average number of multicellular clones//intestine in indicated genotypes and time points.

Interestingly, we also identified cells co-expressing ISC/EE markers in the wild-type tissue near 15% of RPPA clones and 25% of RPMA clones, but not inside the clones themselves (Figure 3d-f). Our results with additional markers are summarized in Figure 3f. For instance, cells coexpressing EB/EE markers were present within and surrounding RPA and RPMA clones and cells coexpressing EB/EC markers within and surrounding RPPA and RPMA clones (Figure S2). We did not observe cells that coexpress multiple cell fate markers in control clones except for rare cells that were positive for both EB and EC markers (Figure 3f). EBs are EC progenitors, so these cells may reflect a transitional stage in EC differentiation. Lastly, we did not observe cells co-expressing ISC/EC markers within or surrounding transformed clones.

The presence of cells that co-express differentiation and stem/progenitor markers within and surrounding transformed clones (Figure 3g) demonstrates that cancer-driving genetic alterations disrupt normal stem cell regulation and differentiation mechanisms. These cells may be products of an abnormal differentiation process resulting in partially differentiated cells that retain some stem cell characteristics. It is also possible that transformed cells exist in more dynamic and fluid states where they can transdifferentiate or dedifferentiate to more stem-like states. To further explore these questions, we focused on the EE cell lineage, which was most frequently affected in our clones. To determine if we can revert fully differentiated EE cells into a proliferative, more stem-like state, we targeted our multigenic combinations to individual EE cells using *pros-gal4* and the PS design. Individual EEs targeted with GFP only never produced multicellular clones, but a significant fraction of those targeted with RPA and RPPA did (Figure 3g-i). Even though multicellular clones derived from EEs were much smaller and did not survive as long (Figure 3g-i), these findings demonstrate that cancer-driving genetic alterations can confer differentiated EEs the ability to proliferate.

### A Context-Dependent Tumor Promoting Role for Enteroendocrine Cells

Our analysis of cell type composition of transformed RPA, RPPA, and RPMA clones demonstrated a significant increase in the relative abundance of the EE cell lineage over time, particularly in smaller clones (Figure 2i,j). Interestingly, human colon tumors with high numbers of EE cells have been associated with more aggressive tumorigenesis and worse prognosis^55,56^, suggesting a tumor-promoting role for EEs. To investigate whether they contribute to the growth of transformed clones, we attempted to deplete EE cells in our clones by knocking down *scute* (*sc*), a transcription factor required for EE differentiation in the adult Drosophila intestine^59^. *sc* knockdown substantially reduced EE cell number within control, RPPA, and RPMA clones but did not affect EE cells within RPA clones (Figure 4a-e). Pros+ EE cells persisted in RPA clones and continued to express the peptide hormone Tk (Figure 4b-e). We next analyzed whether EE depletion affected the growth of transformed clones. We observed a significant reduction in the size of RPMA clones upon EE depletion (Figure 4f). Consistent with the continued presence of the EE cells in RPA clones, *sc* knock-down did not affect the size of RPA clones. Surprisingly, we also found no significant difference in RPPA clone size, despite a significant reduction in EE cell numbers (Figure 4a,f). Interestingly, we often found GFP-negative, wild-type EE cells expressing pros and Tk trapped inside large, EE-depleted RPPA clones (figure 4g,h). We did not observe such cells in the absence of *sc* knock-down in RPPA or any other genotype.

**Figure 4:**
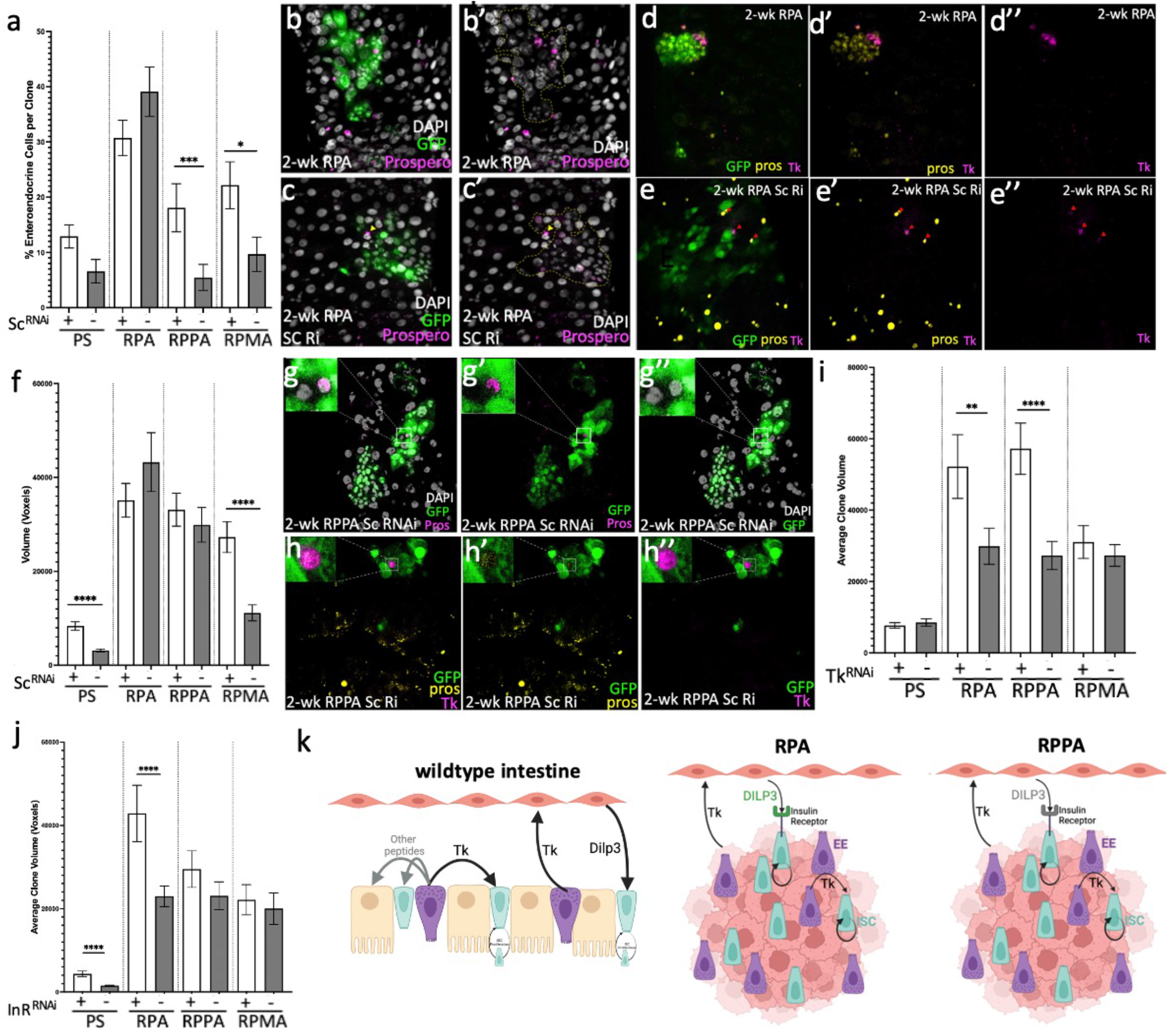
A context-dependent tumor-promoting role for enteroendocrine cells. **(a)** Fraction of pros+ EE cells upon *sc* knock-down in control and transformed clones of different genotypes two weeks post-induction. **(b,c)** Example RPA clones with or without *sc* knock-down two weeks post-induction. Clones are in green, nuclei (DAPI) in gray, and pros in magenta. Yellow arrowhead: a wild-type EE just outside the clone. **(d,e)** Tk expression in RPA clones with or without *sc* knock-down two weeks postinduction. Clones are in green, pros in yellow, and Tk in magenta. Red arrows indicate cells in the clone that are both Pros and Tk positive. **(f)** Volume analysis of control and transformed clones of different genotypes with or without *sc* knock-down two weeks post-induction. **(g)** a GFP negative, Pros+ (magenta) wildtype EE trapped within an EE-depleted RPPA clone. Nuclei (DAPI) are in gray. **(h)** a GFP negative, Pros+ (yellow) wildtype EE expressing Tk (magenta) trapped within an EE-depleted RPPA clone. **(g,h)** The top left corner of each panel shows a magnified view of the GFP-negative cell. **(i,j)** Volume analysis of control and transformed clones of different genotypes with or without *Tk* **(i)** or InR **(j)** knock-down two weeks post-induction. **(k)** A visual depiction of context-dependent contributions of transformed EEs to clone growth (made with BioRender). EE cells support normal intestinal homeostasis by secreting hormone peptides such as Tk to regulate ISC proliferation directly and indirectly. Transformed clones with high numbers of transformed EEs appropriate these mechanisms to support their growth. The specific combination of cancer-driving genetic alterations transformed cells carry determines their dependence on these niche signals. **(a,f,i,j)** Error bars represent the standard error of the mean (SEM). *: P ≤0.05, **: P≤ 0.01, ***: P≤ 0.001, ****: P≤ 0.0001 (Multiple unpaired t-tests with Holm-Šídák correction, PRISM Software)

Enteroendocrine cells serve a niche function to regulate intestinal homeostasis by secreting signals directed to other cell types, including ISCs, both directly and indirectly^34,60,61^. Our findings suggest that depleting EE cells from transformed clones compromises clone growth by depriving transformed clones of these niche signals. Furthermore, at least in the case of RPPA, transformed clones can adapt to this loss by appropriating niche signals secreted by wild-type EE cells. It is unclear whether EE-depleted RPPA clones actively recruit and trap wild-type EE cells or if they are only able to grow if they emerge close to wild-type EE cells. Either way, these findings emphasize the ability of transformed cells to adapt to changes in their cell type composition by diverting niche signals from their environment to support their growth.

A well-established mechanism by which EE cells regulate intestinal homeostasis is through the peptide hormone Tk^32,33^. Tk signals the overlying visceral muscle to express and secrete the Insulin-like peptide3 (DILP3), which is thought to stimulate ISC proliferation through the Insulin Receptor (InR)^32^. We next tested whether transformed EEs contribute to clone growth through this feedback mechanism. Tk knock-down within transformed clones strongly reduced the growth of RPA and RPPA clones (Figure 4i), demonstrating a role for Tk expressed within transformed clones in clone growth. Knocking down InrR, which is thought to mediate ISCs’ response to this feedback mechanism, resulted in a significant reduction in RPA clone size only (Figure 4j). Even though Tk has a critical role in promoting the growth of both RPA and RPPA clones, RPPA appears to be insensitive to the specific feedback mechanism involving Tk/DILP3/InR axis, likely because both MAPK and PI3K pathways are already strongly activated in this model due to PTEN loss. Tk must contribute to RPPA clone growth by signaling other cell types in the intestine. For instance, Tk can signal ECs to regulate lipid metabolism through its receptor TkR99D; whether Tk-TkR99D signaling directly regulates ISC proliferation remains to be determined^34,62^.

Furthermore, even though transformed EEs play an essential role in the growth of RPMA clones (Figure 4f), this role appears to be independent of Tk, as these clones are not sensitive to Tk or InR knock-down (Figure 4i,j). Combined, these studies demonstrate that the specific combination of genetic alterations in a transformed cell impacts its lineage identity and determines whether and how it responds to niche signals in its environment (Figure 4k). Understanding these context-dependent nuances and their impact during intestinal transformation could identify novel vulnerabilities and druggable regulatory nodes that could be exploited for targeted therapy in specific genetic contexts.

## DISCUSSION

Here, we report a new genetic platform, PromoterSwitch, designed to target transgene expression to a small subset of cells expressing any tissue/cell-type specific Gal4 of interest. We used this platform to generate and study transformed clones by lineage-specific genetic manipulations targeted to a small number of individual intestinal cells. By targeting fewer cells in each intestine, we were able to generate and study the behavior of discrete clones over time without compromising tissue integrity and survival. As the initially targeted cells and their progeny differentiated into other cell types, they continued to express our transgene combinations, allowing us to establish clones that captured the cell type heterogeneity of the normal intestinal epithelium and human colon tumors. Consolidating the transgenes necessary for this complex experimental platform using our multigenic vectors^63^ simplified the genetic background of our experimental animals, allowing additional genetic manipulations for mechanistic studies.

PromoterSwitch is a flexible platform that can be combined with any gal4 of interest to achieve more refined and permanent genetic manipulations. For instance, it could be used to reduce the number of targeted cells in other disease models to better study their interactions with their environment. It will also be particularly effective for stem cell biology and developmental biology studies where cells of interest often shut down cell-type/tissue-specific Gal4 expression as part of their developmental and differentiation program.

Like most epithelial tumors, colorectal cancer is a genetically complex and heterogeneous disease where concurrent deregulation of multiple cancer-relevant pathways is common. Our RPA model, which combines APC and TP53 loss with oncogenic KRAS, represents one of the most common genomic landscapes in colorectal cancer. However, most colon tumors include deregulation of additional pathways, most commonly PI3K and TGF-β pathways, respectively represented by PTEN loss and SMAD4 (Medea) loss in our RPPA and RPMA models. Our analysis revealed significant differences in proliferation rates and growth trajectories of transformed clones with different genotypes, emphasizing the importance of the specific genomic landscape of a transformed cell in determining its intrinsic properties and behavior.

Previous research has shown that most tumors retain the cell-type heterogeneity found in their tissue of origin and that intrinsic, lineage-specific differences between cell types can affect tumor progression, evolution, prognosis, and treatment response^55,64–66^. Our transformed clones also captured the cell lineage heterogeneity of human colon tumors. In addition, we found differences in the relative abundance of different cell lineages, indicating disruptions in stem cell renewal, differentiation, and cellcell communication mechanisms among different intestinal cell types. For instance, we found a significant enrichment of the EE lineage in all multigenic models and noted that larger clones appeared to be more undifferentiated.

In addition to the changes in the relative abundance of different cell lineages in our transformed clones, we also identified novel mixed cell lineages characterized by the co-expression of two intestinal cell type markers. While the specific markers that were co-expressed vary by the genotype of the model, the presence of EE cells that co-express stem and progenitor cell markers are the most common. The presence of Dl-positive cells that co-express EE-specific peptide hormone Tk and the ability of differentiated EEs to give rise to small, multicellular clones suggest that transformed clones have cells that can perform functions associated with multiple lineages, highlighting the dynamic and highly plastic nature of transformed cells.

Stem cell function and, more broadly, epithelial homeostasis is regulated by complex interactions that integrate short and longer-range signals with intrinsic, lineage-specific properties of individual cells. These already complex intercellular signaling mechanisms are then disrupted by cancer-driving genetic alterations in context-dependent and often unpredictable ways and appropriated to support tumor growth. This point is illustrated by the genotype-dependent nuances in molecular mechanisms underlying the tumor-promoting role we uncovered for the EE lineage in our multigenic models.

We found that both RPA and RPPA clones are dependent on the peptide hormone Tk secreted from transformed EEs to promote their growth, but the mechanisms appear to be different: The Tk/Dilp3/InR axis, a feedback mechanism by which EEs regulate ISC proliferation via the overlying visceral muscle, appears to have an essential role in supporting the growth of RPA clones. On the other hand, RPPA is not dependent on this feedback mechanism, suggesting that different Tk-mediated cell communication pathways play a role. RPMA clones, while dependent on transformed EEs to support their growth, are not sensitive to Tk loss, suggesting that the tumor-promoting function of EEs in this particular genetic context is mediated by other signals secreted by the transformed EEs^33,34,67^. Furthermore, our observation that the EE cell fate in RPA clones doesn’t appear to require the transcription factor Scute, which is essential for the EE cell fate in the adult intestine^59,^, highlights genotype-dependent differences in EE cell identity in our models. Lastly, our observation that EE-depleted RPPR clones can appropriate non-transformed, wild-type EEs to support their growth demonstrates the highly adaptable nature of transformed clones.

## MATERIALS AND METHODS

### Cloning of PromoterSwitch Construct and transgenesis

PromotorSwitch transgene was first digitally assembled using publicly available sequences. Two pieces, [FRT-FRT3-Gal80^ts^-FRT] and [Gal80-FRT3], were generated by gene synthesis (Genewiz/Azenta). act5C promoter and Gal4 coding sequences were amplified from the CoinFLP plasmid (Addgene 52890)^54^ as two separate fragments. All four fragments were sequentially cloned into pUC57-Kan by standard restriction cloning methods to generate the final [Act5C-FRT-FRT3-Gal80^ts^-FRT-Gal80-FRT3] transgene.

To generate the multigenic PS construct that included the [UAS-GFP], [UAS-FLP], and the [Act5C-FRT-FRT3-Gal80^ts^-FRT-Gal80-FRT3] transgenes, we modified our multigenic vector^53^ by removing one of the three UAS cassettes and cloning FLP and GFP coding sequences into the multiple cloning sites of the remaining two UAS cassettes. FLP and GFP coding sequences used for cloning were first PCR-amplified from genomic DNA of transgenic flies carrying these transgenes using primers designed to append restriction sites of enzymes NotI and XbaI to the 5’ and 3’ end of the FLP recombinase coding sequence, and BsiWI and AsiSI to the 5’ and 3’ end of the GFP coding sequence, respectively. Lastly, the [Act5C-FRT-FRT3-Gal80^ts^-FRT-Gal80-FRT3] fragment was cloned into the multigenic vector from pUC57-Kan using PmeI and AgeI. Each piece was sequence-confirmed before they were used for cloning,

After the cloning process was completed, all three transgenes in the final multigenic PromoterSwitch construct were sequence-confirmed, and transgenic flies were generated by PhiC31-mediated targeted integration into the *attp40* and *attp2* landing sites on the second and third Drosophila chromosomes, respectively. Both second and third chromosome insertions were functionally validated; for this work, we used one of the third chromosome insertions (attp2, line M2)

### Drosophila Strains

All strains were maintained at room temperature on a standard Drosophila medium. Multigenic cancer combinations used in this study are described previously^50^. Transgenic Drosophila lines used in multigenic cancer combinations are *UAS-ras^G12V^* (II, G. Halder), and three RNAi lines, *UAS-p53^RNAi^* (II), *UAS-pten^RNAi^* (III) and *UAS-apc^RNAi^* (II), that were obtained from the Vienna Drosophila Resource Center VDRC^68^. Additional transgenic Drosophila lines used in this study were obtained from the Bloomington Drosophila Stock Center (BDSC): *Dl-LacZ* (BDSC #11651), *Su(H)GBE-LacZ* (BDSC #83352), UAS-*sc^shRNA^* BDSC #26206), *UAS-Tk^RNAi^ (*BDSC #25800) and *UAS-InR^RNAi^* (BDSC # 35251).

A third chromosome insertion of the PS construct (line M2, inserted into the attp2 landing site) was incorporated into the background of each cancer model to generate the following fly lines: **1)** *w^1118^; UAS-apc^RNAi^/CyO; PS attp2 M2/TM6b, Hu, Tb* **2)** *w^1118^; UAS-ras^G12V^ UAS-p53^RNAi^ UAS-apc^RNAi^/CyO; PS attp2 M2/TM6b, Hu, Tb* **3)** *w^1118^; UAS-ras^G12V^ UAS-p53^RNAi^ UAS-pten^RNAi^ UAS-apc^RNAi^/7CyO; PS attp2 M2/TM6b, Hu, Tb* and **4)** *w^1118^; UAS-ras^G12V^ UAS-p53^RNAi^ UAS-pten^RNAi^ UAS-Med^RNAi^/CyO; PS attp2 M2/TM6b, Hu, Tb*.

Experimental animals were generated by crossing virgin females from the *w^1118^ UASdcr-2/Y, hs-hid; esg-gal4 tub-gal80^ts^* or *w^1118^ UASdcr-2/Y, hs-hid; pros-gal4/TM6b, Hu, Tb* to males carrying multigenic combinations and the PS construct listed above. Males carrying the PS construct only were used as controls. The Y chromosome hs-hid transgene, which results in ubiquitous activation of apoptosis when induced^69^, was used to kill all male progeny and facilitate mass virgin female collection. Crosses were kept at 18°C to prevent transgene expression. When progeny emerged, progeny with the following genotypes were collected: For *esg-gal4:* **1)** *w^1118^ UASdcr-2/w^1118^; UAS-apc^RNAi^/esg-gal4 tub-gal80^ts^; PS attp2 M2/+* **2)** *w^1118^ UASdcr-2/w^1118^; UAS-ras^G12V^ UAS-p53^RNAi^ UAS-apc^RNAi^/esg-gal4 tub-gal80^ts^; PS attp2 M2/+* **3)** *w^1118^ UASdcr-2/w^1118^; UAS-ras^G12V^ UAS-p53^RNAi^ UAS-pten^RNAi^ UAS-apc^RNAi^/esg-gal4 tub-gal80^s^; PS attp2 M2/+* **4)** *w^1118^ UASdcr-2/w^1118^; UAS-ras^G12V^ UAS-p53^RNAi^ UAS-pten^RNAi^ UAS-Med^RNAi^/esg-gal4 tub-gal80^ts^; PS attp2 M2/+* and **5)** *w^1118^ UASdcr-2/w^1118^; +/esg-gal4 tub-gal80^ts^; PS attp2 M2/+* as controls. For *pros-gal4:* **1)** *w^1118^ UASdcr-2/w^1118^; UAS-ras^G12V^ UAS-p53^RNAi^ UAS-apc^RNAi^/+; PS attp2 M2/pros-gal4* **2)** *w^1118^ UASdcr-2/w^1118^; UAS-ras^G12V^ UAS-p53^RNAi^ UAS-pten^RNAi^ UAS-apc^RNAi^/+; PS attp2 M2/pros-gal4* and **5)** *w^1118^ UASdcr-2/w^1118^; +/+; PS attp2 M2/pros-gal4* as controls. Clones were induced by placing collected flies at 29°C to inactivate Gal80^ts^. Experimental animals were transferred onto fresh food three times a week until dissection.

To introduce the Lac-Z and RNAi transgenes into experimental animals, the following fly lines were generated and crossed to males with genotypes listed above: **1)** *w^1118^ UASdcr-2/Y, hs-hid; esg-gal4 tub-gal80^ts^; Dl-LacZ /TM6b, Hu, Tb* **2)** *w^1118^ UASdcr-2/Y, hs-hid; esg-gal4 tub-gal80^ts^/CyO; Su(H)-GBE-LacZ /TM6b, Hu, Tb* **3)** *w^1118^ UASdcr-2/Y, hs-hid; esg-gal4 tub-gal80^ts^; UAS-sc^shRNA^/S-T, tub-gal80, Cy, Tb, Hu* **4)** *w^1118^ UASdcr-2/Y, hs-hid; esg-gal4 tub-gal80^ts^; UAS-Tk^RNAi^/S-T, tub-gal80, Cy, Tb, Hu* and **5)** *w^1118^ UASdcr-2/Y, hs-hid; esg-gal4 tub-gal80^ts^; UAS-InR^RNAi^/S-T, tub-gal80, Cy, Tb, Hu*

### Immunohistochemistry, Imaging, and Scoring

Adult female *Drosophila* intestines were dissected in Phosphate Buffered Saline (PBS) and fixed for 15 minutes at room temperature in ice-cold 4% paraformaldehyde in PBS. Intestines were washed in PBS and blocked in PBGT (PBS containing 0.1% Triton X and 1% normal goat serum) for 1 hour at room temperature, incubated overnight with primary antibodies at 4°C, rinsed in PBS three times, followed by a 1-hour block at room temperature and incubated with secondary antibody for 2 hours at room temperature. Intestines were mounted in VectaShield mounting medium containing DAPI.

Primary antibodies used were; mouse anti-phospho-SAPK/JNK-pThr183/pTyr185 G9 (Cell Signaling Technology, #9255), Rabbit anti-phospho-SRC-pTyr419 (Thermo Fisher Scientific, 44-660-G), rabbit anti-phospho-Histone-H3-pSer10 (Sigma Aldrich, H0412), rabbit anti-phospho-AKT-pSer505 (Cell Signaling Technology, #4054), mouse anti-diphospho-ERK1/2 (Sigma Aldrich, M8159), mouse anti-MMP1 (DSHB, 3B8D12), rabbit anti-beta galactosidase (Thermo Fisher Scientific, # A-11132), mouse anti-Prospero (DSHB, MR1A), mouse anti-Nubbin (DSHB, 2D4), and rabbit anti-Tachykinin (a generous gift from Dr. Jan A. Veenstra, The University of Bordeaux). Alexa 568- or 633-conjugated goat-anti-mouse and goat-anti-rabbit antibodies were used as secondary antibodies.

Fluorescence images were captured using a Leica TCS SPE-II DM6 confocal microscope under a 40X objective and processed using Leica LAS-AF software. Additional processing required for quantification were performed using Fiji Image J Software^70^. Approximately 20 flies were dissected for each genotype, and over 100 clones were analyzed for each experiment. Results were confirmed using an independent set of experiments.

### Quantification of clone size and cell lineage composition using CellProfiler

Images were quantified for clone size and the number of ISCs, EBs, EEs, and ECs per clone (separately) using CellProfiler cell image analysis software^71,72^. Customized pipelines designed for this analysis are available upon request.

Briefly, GFP+ clones and DAPI+ cells were identified using the MedianFilter algorithm. DAPI and GFP segmentations were overlaid using the OverlayObjects module, and clone sizes were quantified in voxels using the MeasureObjectSizeShape module. The number of nuclei per clone was quantified using the RelateObjects algorithm and the output from the overlayobjects module. For analyses including staining for cell lineage markers, the GFP channel, the channel containing the cell lineage markers, and the DAPI channel were segmented using the Watershed algorithm, processed using the OverlayObjects module. Then the RelateObjects algorithm was also used to quantify the number of each cell type per clone. To quantify the non-autonomous pJNK activity, the MedianFilter module was used to identify GFP+ clones and the pJNK signal. The watershed median was used to segment clones and the pJNK signal individually. Next, the MaskImage module was used to mask the GFP channel containing the clones. The OverlayObjects module was then used on the mask and pJNK images, and the MeasureObjectSizeShape module was used to quantify in voxels the volume of the pJNK channel in GFP-cells surrounding the clones.

## SUPPLEMENTARY FIGURE LEGENDS

**Supplementary Figure 1.**
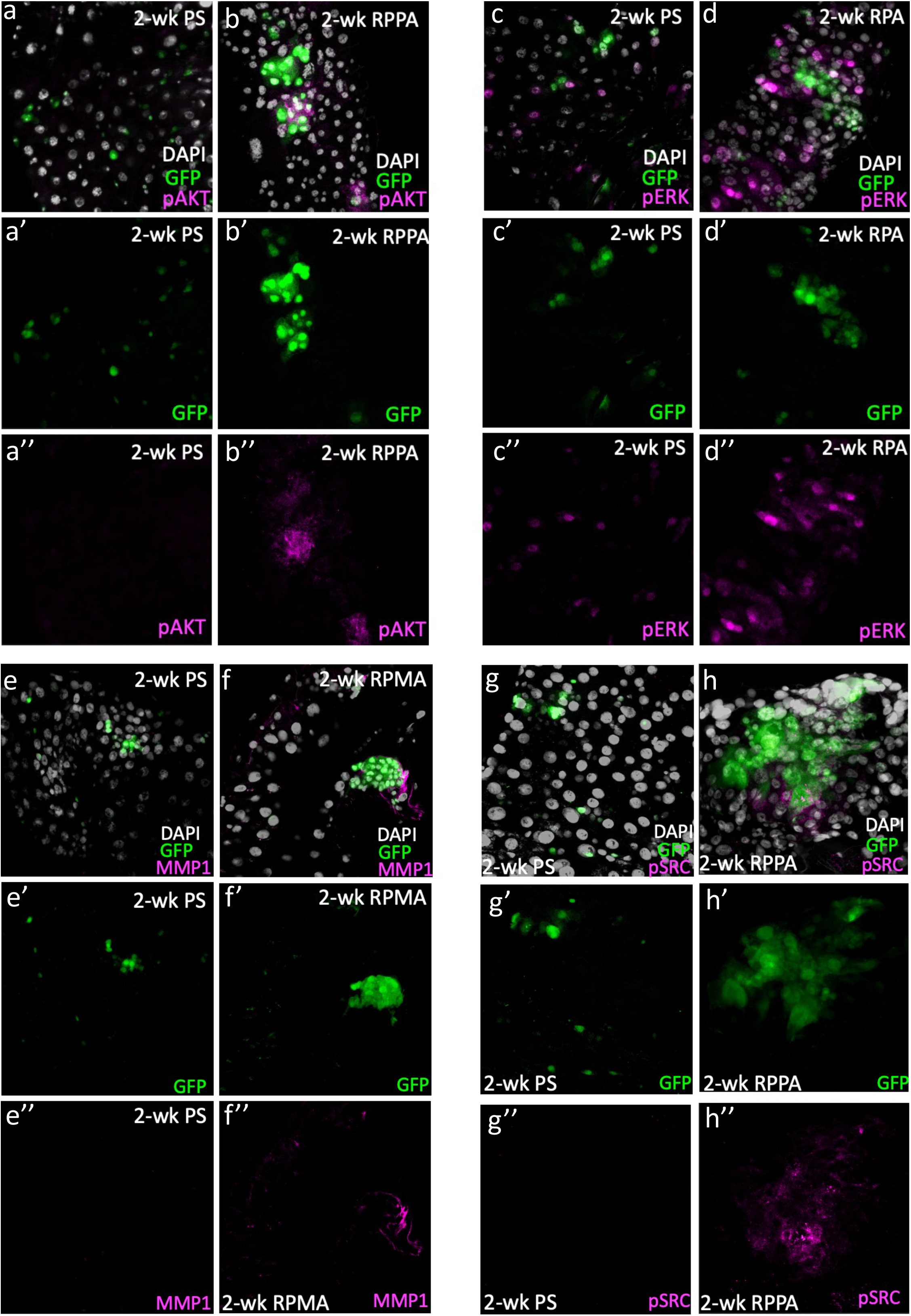
Activation of cancer-relevant pathways within and surrounding transformed clones. **(a-b)** phospho-AKT (pAKT, in magenta) staining in intestines with two-week-old control **(a)** and RPPA **(b)** clones (green). **(c-d)** phospho-ERK (pERK, in magenta) staining in intestines with two-week-old control **(c)** and RPA **(d)** clones (green). **(e-f)** MMP1 (in magenta) staining in intestines with two-week-old control **(e)** and RPMA **(f)** clones (green). **(g-h)** phospho-SRC (pSRC, in magenta) stainings in intestines with two-week-old control **(g)** and RPPA **(h)** clones (green). Intestines carrying transformed clones of all genotypes show both autonomous and non-autonomous activation of each marker. A single example for each marker and its matching control is presented for brevity.

**Supplementary Figure 2.**
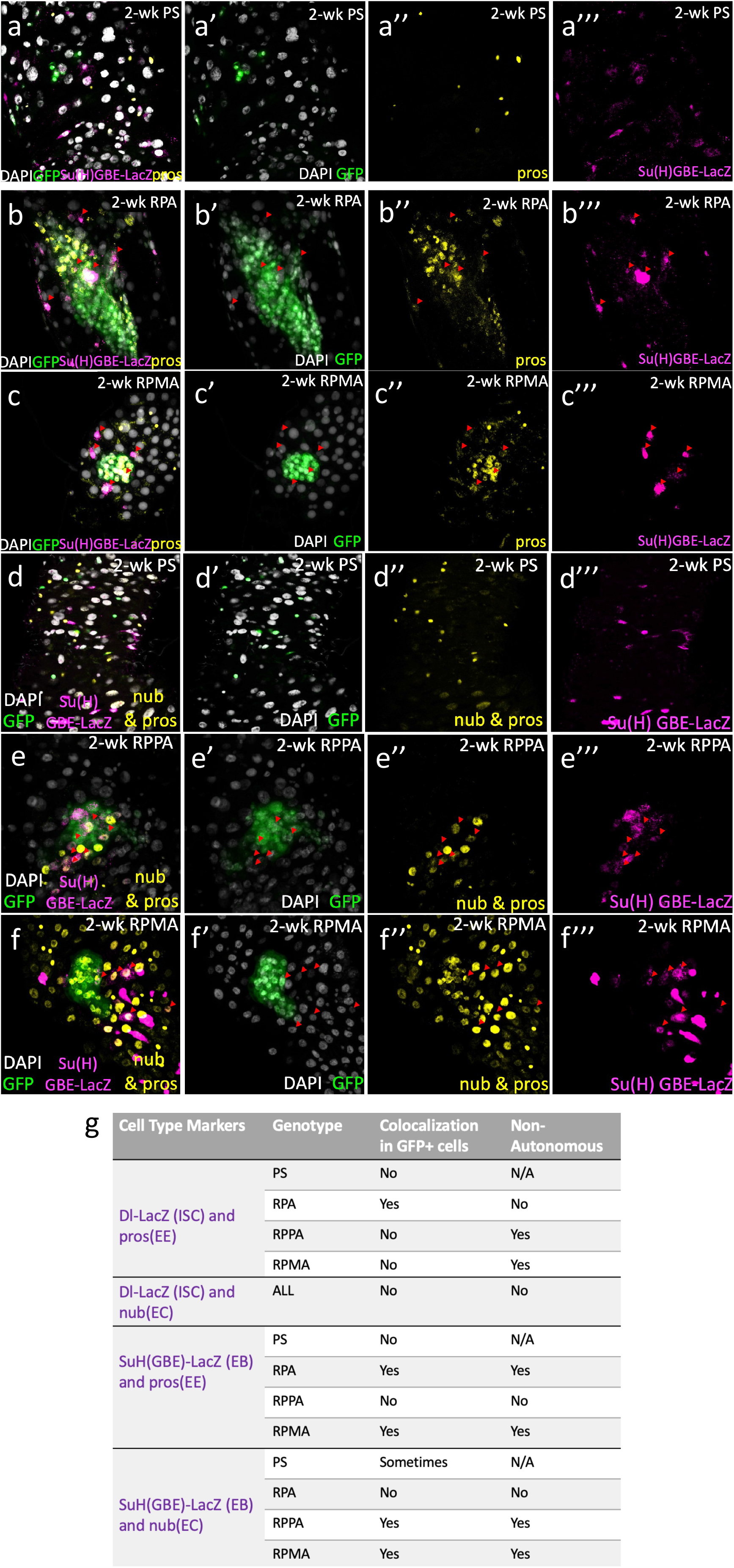
Transformed clones contain cells that co-express multiple cell fate markers. **(a-c)** Intestines with example GFP-only control **(a)**, RPA **(b),** and RPMA **(c)** clones, immunostained for indicated markers two weeks after induction. Clones are in green, nuclei (DAPI) in gray, pros in yellow, Su(H) GBE-LacZ in magenta. **(d-f)** Intestines with GFP-only control **(d)**, RPPA **(e),** and RPMA **(f)** clones, immunostained for indicated markers two weeks after induction. Clones are in green, nuclei (DAPI) in gray, nubbin and pros in yellow, and Su(H)GBE-LacZ in magenta. Red arrows pointing towards the bottom left of images indicate cells co-expressing lineage markers. **(g)** Table summarizing mixed lineages observed within and surrounding clones with different genotypes.

## ACKNOWLEDGEMENTS

This study used transgenic RNAi lines (Office of the Director R24 OD030002: “TRiP resources for modeling human disease”) obtained from the Bloomington Drosophila Stock Center (NIH P40OD018537) and the Vienna Drosophila Resource Center (VDRC, www.vdrc.at), and monoclonal antibodies obtained from the Developmental Studies Hybridoma Bank, created by the NICHD of the NIH and maintained at The University of Iowa, Department of Biology, Iowa City, IA 52242. We thank Alexander Teague for essential discussions and technical assistance during the early stages of this work. We thank Jason Cassara, Justin Brown, Eric Cruz, Sneha Kapil, Autumn Hawkins, and Carson Caddy for technical support. This work was supported by grants from the National Institutes of Health (R03 CA219321 and R21 GM141734).

## Notes

### Competing Interest Statement

The authors have declared no competing interest.

